# Taxonomic repositioning of twelve *Xanthomonas campestris*, seven *Xanthomonas axonopodis* and one *Pseudomonas cissicola* reference pathovars to *Xanthomonas citri*

**DOI:** 10.1101/2021.07.23.453582

**Authors:** Kanika Bansal, Sanjeet Kumar, Prabhu B. Patil

**Affiliations:** CSIR- Institute of Microbial Technology, Chandigarh, India

**Keywords:** *Xanthomonas citri* pathovars, reference pathovars, taxonogenomics, phylogenomics

## Abstract

Here on the basis of phylo-taxonogenomics criteria we present amended description of 20 pathovars to *Xanthomonas citri* majority (18/20) of which were first reported from India. 7/20 are currently classified as *X. axonopodis*, 12/20 as *X. campestris* and 1/20 as *Pseudomonas cissicola*. Here, we have generated genome sequence data for 4 pathovars and genomes of remaining 16 were used from the published data. Genome based investigation reveals that all these pathovars belong to *X. citri* and not to *X. axonopodis* or *X. campestris* as previously reported. Present proposal is to aid in resolving the taxonomic confusion of the *X. ctiri* pathovars and prevent future usage of invalid names.

---

*Xanthomonas* sp. bears high degree of phenotypic and 16S rRNA similarity amongst diverse range of phytopathogens (Hayward 1993, Hauben, Vauterin et al. 1997, Moore, Krüger et al. 1997). Consequently, classical approach of taxonomy has resulted in various ambiguities in its classification and various revisions in the taxonomy of the constituent species (Vauterin, Rademaker et al. 2000, Naushad, Adeolu et al. 2015, Kumar, Bansal et al. 2019, Bansal, Kumar et al. 2020). *Citrus spp*. is associated with a vast range of *Xanthomonas* strains following pathogenic and non-pathogenic lifestyles (Hasse 1916, Vauterin, Yang et al. 1996, Bansal, Kumar et al. 2020). Genome-based approaches deciphered these non-pathogenic strains to be closely related to *X. sontii* which were earlier not resolved by classical taxonomy (Bansal, Kaur et al. 2019, Bansal, Midha et al. 2019). On the other hand, the pathogenic counterpart *X. citri* pv. citri is known over a century as the causal agent of citrus canker having a quarantine status (Raychaudhuri, Verma et al. 1972, Gottwald, Graham et al. 2002, Brunings and Gabriel 2003, Graham, Gottwald et al. 2004, Gottwald and Irey 2007). Earlier it was classified as *X. campestris* pv. citri (Group A) (ex Hasse, 1915) and then as *X. axonopodis* pv. citri based on classical approaches like: restriction fragment length polymorphism (RFLP), DNA-DNA hybridization etc. (Gabriel, Kingsley et al. 1989, Vauterin, Hoste et al. 1995). In the last two decades, numerous attempts were made to resolve the taxonomy and phylogeny of the strains belonging to *X. citri* (Schaad, Postnikova et al. 2006, Ah-You, Gagnevin et al. 2009) resulting in various revisions in taxonomy.

Detailed genomic investigation of the *X. axonopodis* species revealed that strains previously classified as *X. axonopodis* instead belong to four different species. These four species were namely, *X. citri, X. axonopodis, X. euvesicatoria* and *X. phaseoli* forming four distinct phylogroups (Constantin, Cleenwerck et al. 2016). Constantin and coworkers have identified fourteen different pathovars isolated form diverse host range constituting the *X. citri* species at the whole genome level. However, besides these pathovars of *X. citri* there are reports of twenty other reference pathovars majorly isolated from India over the century from diverse commercially important hosts (Parkinson, Aritua et al. 2007, Parkinson, Cowie et al. 2009). Due to limitations of classical taxonomic methods, these were classified as different species of *Xanthomonas* like *X. campestris, X. axonopodis* and one was even classified as *Pseudomonas cissicola* (Takimoto 1939). However, *gyrB* based phylogeny have revealed their phylogenetic relatedness with the *X. citri* (Parkinson, Aritua et al. 2007, Parkinson, Cowie et al. 2009). Further, whole genome level investigation of these reference pathovars also suggests the need of revisiting their taxonomy (Bansal, Midha et al. 2017).

These reference pathovars include *X. campestris* pv. durantae isolated from *Duranta repens* which is clonal to the *X. citri* pv. citri at the whole genome level (Bansal, Midha et al. 2017). Similarly, *X. campestris* pv. centellae isolated from *Centella asiatica* is clonal to *X. campestris* pv. mangiferaeindicae causing mango bacterial black spot leading substantial loss in fruit quality in Asia, Africa, Central America, Caribbean and Oceania (https://www.cabi.org/ISC/abstract/20123367489); (Pruvost 1993, Pruvost, Couteau et al. 1998). Additionally, 18 pathovars isolated from diverse hosts namely, *Cissus woodrowii, Bauhinia racemosa, Martynia diandra, Cayratia trifolia, Khaya senegalensis, Cayratia japonica, Tectona grandis, Aegle marmelos, Azadirachta indica, Cajanus cajan, Clitoria* sp., *Thespesia populnea* and *Leea asiatica, Sesbania aegyptiaca, Merremia gangetica, Triumfetta pilosa* and *Trichodesma zeylanicum* are included in the present study (table 1). Interestingly, all of these reference pathovars were first reported from India except for two pathogens i.e., *X. axonopodis* pv. khayae LMG 753 and *Pseudomonas cissicola* LMG 21719 reported from Sudan and Japan respectively (table 1). There is confusion surrounding the taxonomy of these reference pathovars. Genome-based investigation of these *X. citri* pathovars (XCPs) have revealed their phylogenetic relationship with the *X. ctiri* (Bansal, Midha et al. 2017).

**Table 1:** Metadata of the strains used in the present study. Strains sequenced in the preset study are highlighted in green color.

| S. No. | Pathovar | Accession no. | Isolation Year | Country | Host | Host Taxonomy | Reference |
| --- | --- | --- | --- | --- | --- | --- | --- |
| 1. | *X. campestris* pv. *vitiswoodrowii* LMG 954 | LOKG00000000 | 1961 | India | *Cissus woodrowii* | Vitaceae, Vitales, Rosids, Eudicots | (Patel and Kulkarni 1951a) |
| 2. | *X. axonopodis* pv. *bauhiniae* LMG 548 | LOKR00000000 | 1961 | India | *Bauhinia racemosa* | Fabaceae, Fabales, Rosids, Eudicots | (Padhya, Patel et al. 1965a) |
| 3. | *X. axonopodis* pv. *martyniicola* LMG 9049 | LOJX00000000 | 1958 | India | *Martynia diandra* | Martyniaceae, Lamiales, Asterids, Eudicots | (Moniz and Patel 1958) |
| 4. | *X. campestris* pv. *vitiscarnosae* LMG 939 | LOKI00000000 | 1962 | India | *Cayratia trifolia* | Vitaceae, Vitales, Rosids, Eudicots | (Moniz and Patel 1958) |
| 5. | *X. campestris* pv. *vitistrifoliae* LMG 940 | LOKH00000000 | 1961 | India | *Cayratia trifolia* | Vitaceae, Vitales, Rosids, Eudicots | (Padhya, Patel et al. 1965b) |
| 6. | *X. axonopodis* pv. *khayae* LMG 753 | LOKN00000000 | 1957 | Sudan | *Khayasenegalensis* | Meliaceae, Sapindales, Rosids, Eudicots | (Sabet 1959) |
| 7. | *Pseudomonas cissicola* LMG 21719 | LOJT00000000 | 1974 | Japan | *Cayratia japonica* | Vitaceae, Vitales, Rosids, Eudicots | (Takimoto 1939) |
| 8. | *X. axonopodis* pv. *melhusii* LMG 9050 | LOJW00000000 | 1961 | India | *Tectonagrandis* | Lamiaceae, Lamiales, Asterids, Eudicots | (Patel, Kulkarni et al. 1952b) |
| 9. | *X. campestris* pv. *bilvae* NCPPB 3213 | CDHI01000000 | 1982 | India | *Aegle marmelos* | Rutaceae, Sapindales, Rosids, Eudicots | (Chakravarti, Sarma et al. 1984) |
| 10. | *X. campestris* pv. *azadirachtae* LMG 543 | LOKS00000000 | 1971 | India | *Azadirachta indica* | Meliaceae, Sapindales, Rosids, Eudicots | (Desai, Gandhi et al. 1966) |
| 11. | *X. campestris* pv. *durantae* LMG 696 | LOKP00000000 | 1956 | India | *Durantarepens* | Verbenaceae, Lamiales, Asterids, Eudicots | (Srinivasan and Patel 1957) |
| 12. | *X. citri* pv. *citri* LMG 9322 | MDJT00000000 | 1915 | United States | *Citrus aurantifolia* | Rutaceae, Sapindales, Rosids, Eudicots | (Hasse 1916) |
| 13. | *X. axonopodis* pv. *cajani* LMG 558 | LOKQ00000000 | 1950 | India | *Cajanuscajan* | Fabaceae, Fabales, Rosids, Eudicots | (Kulkarni 1950) |
| 14. | *X. axonopodis* pv. *clitoriae* LMG 9045 | LOKA00000000 | 1974 | India | *Clitoria* sp. | Fabaceae, Fabales, Rosids, Eudicots | (Pandit and Kulkarni 1979) |
| 15. | *X. campestris* pv. *centellae* LMG 9044 | LOJR00000000 | 1979 | India | *Centella asiatica* | Apiaceae, Apiales, Asterids, Eudicots | (Basnyat and Kulkarni 1979) |
| 16. | *X. campestris* pv. *thespesiae* LMG 9057 | LOJU00000000 | 1978 | India | *Thespesia populnea* | Malvaceae, Malvales, Rosids, Eudicots | (Patil and Kulkarni 1981) |
| 17. | *X. campestris* pv. *leeana* LMG 9048 | LOJY00000000 | 1967 | India | *Leea asiatica* | Vitaceae, Vitales, Rosids, Eudicots | (Patel and Kotasthane 1969a) |
| 18. | *X. axonopodis* pv. *sesbaniae* NCPPB 582 | LOKL00000000 | 1958 | India | *Sesbania aegyptiaca* | Fabaceae, Fabales, Rosids | (Patel, Kulkarni et al. 1952) |
| 19. | *X. campestris* pv. *merremiae* NCPPB 3114 | LOJV00000000 | 1976 | India | *Merremia gangetica* | Convolvulaceae, Solanales, Asterids | (Pant and Kulkarni 1978) |
| 20. | *X. campestris* pv. *thirumalacharii*<br>NCPB 1452 | LOKK0000000<br>0 | 1961 | India | *Triumfetta pilosa* | Malvaceae, Malvales, Rosids | (Padhya and Patel<br>1964) |
| 21 | *X. campestris* pv. *trichodesmae* NCPB<br>585 | LOKJ00000000 | 1953 | India | *Trichodesma*<br>*zeylanicum* | Boraginaceae, Boraginales, Asterids | (Patel, Kulkarni et al.<br>1952b) |

For the present study, genomes of four pathovars namely, *X. axonopodis* pv. sesbaniae NCPPB 582, *X. campestris* pv. merremiae NCPPB 3114, *X. campestris* pv. thirumalacharii NCPPB 1452 and *X. campestris* pb. trichodesmae NCPPB 585 were obtained by Illumina MiSeq sequencing by following method as described by Bansal *et al* (Bansal, Midha et al. 2017) and genome statistics are provided in table 2. Here, genome size of the strains was around 5 MB with approximately 64.6% GC content (table 2).

**Table 2:** Genome statistics of the sequenced strains in the present study.

| S. No. | Pathovar | Genome size (bp) | No. of contigs | Coverage (X) | GC content (%) | N50 (bp) | No. of genes | No. of tRNAs+rRNA+ncRNA | NCBI accession no. |
| --- | --- | --- | --- | --- | --- | --- | --- | --- | --- |
| 1 | *X. axonopodis* pv. *sesbaniae* NCPPB 582 | 54,04,710 | 190 | 140 | 64.52 | 1,04,268 | 4,703 | 52+2+1 | LOKL00000000 |
| 2 | *X. campestris* pv. *merremiae* NCPPB 3114 | 50,34,542 | 135 | 167 | 64.67 | 1,17,975 | 4,215 | 53+3+1 | LOJV00000000 |
| 3 | *X. campestris* pv. *thirumalacharii* NCPPB 1452 | 49,27,392 | 291 | 177 | 64.77 | 42,427 | 4,311 | 50+3+1 | LOKK00000000 |
| 4 | *X. campestris* pv. *trichodesmae* NCPPB 585 | 54,73,142 | 174 | 167 | 64.57 | 1,08,567 | 4,682 | 52+5+1 | LOKJ00000000 |

To access their taxonomic status, various parameters of taxonogenomics like average nucleotide identity (ANI) and digital DNA-DNA hybridization (dDDH) were implemented. ANI and dDDH were calculated using orthoANI v1.2 (Yoon, Ha et al. 2017) and GGDC 2.1 (http://ggdc.dsmz.de/ggdc.php#) respectively. *X. citri* pv, citri LMG 9322 (T) and *X. citri* pv. fuscans LMG 826 (T) were having similar ANI and dDDH values with the reference pathovars i.e., >96% and >67% respectively (table 3). Whereas, reference pathovars were having minimum ANI values of 92%, 84% and dDDH values of 49%, 29% with the type strains *X. axonopodis* DSM 3585 (T) and *X. campestris* pv. campestris ATCC 33913 (T) respectively. Genome similarity assessment clearly demarcated reference pathovars under study (n=20) from *X. axonopodis* and *X. campestris*. Further, ANI values of >96% clearly indicates that all twenty reference pathovars belongs to *X. citri*. However, dDDH values among *X. citri* pv. citri LMG 9322 (T) and *X. citri* pv. fuscans LMG 826 (T) is 68.6% yet they belong to same species i.e. *X. citri* (Constantin, Cleenwerck et al. 2016). In the current study as well the dDDH values for *X. axonopodis* pv. sesbaniae NCPPB 582, *X. campestris* pv. merremiae NCPPB 3114, *X. campestris* pv. thirumalacharii NCPPB 1452 and *X. campestris* pv. trichodesmae NCPPB 585 were in the range of 67.2% - 68.3% and 88% - 89.7% for *X. citri* pv. citri LMG 9322 (T) and *X. citri* pv. fuscans LMG 826 (T) respectively. Hence, genome similarity values with the type strains of *X. citri, X. axonopodis* and *X. campestris* clearly depicted that all the twenty reference pathovars under study belong to *X. citri*.

**Table 3:** Digital DNA-DNA hybridization values of the reference pathovars with the *X. citri* pv. citri LMG 9322 (T), *X. axonopodis* DSM 3585 and *X. campestris* pv. campestris ATCC 33913 (T).

|  | *X. citri* pv. citri LMG 9322 (T) |  | *X. axonopodis* DSM 3585 (T) |  | *X. campestris* pv. campestris ATCC 33913 (T) |  | *X. citri* pv. fuscum LMG 726 (T) |  |
| --- | --- | --- | --- | --- | --- | --- | --- | --- |
|  | dDDH | ANI | dDDH | ANI | dDDH | ANI | dDDH | ANI |
| *X. citri* pv. citri LMG 9322 (T) | 100 | 100 | 50.3 | 93.1 | 30 | 85.1 | 68.6 | 96.3 |
| *X. campestris* pv. durantae LMG 696 | 98.4 | 99.8 | 49.8 | 92.9 | 29.7 | 85.2 | 68 | 96.2 |
| *X. axonopodis* pv. clitoriae LMG 9045 | 92.2 | 99.1 | 49.8 | 93 | 29.5 | 85 | 67.6 | 96.1 |
| *X. axonopodis* pv. cajani LMG 558 | 92.2 | 99.1 | 49.6 | 92.9 | 29.7 | 85.1 | 68.3 | 96.3 |
| *X. campestris* pv. centellae LMG 9044 | 90.8 | 98.8 | 49.6 | 93 | 29.7 | 85.1 | 68.4 | 96.3 |
| *X. campestris* pv. bilvae NCPPB 3213 | 89.9 | 98.8 | 49.6 | 92.9 | 29.8 | 84.9 | 68.5 | 96.2 |
| *X. axonopodis* pv. melhusii LMG 9050 | 89.6 | 98.8 | 49.6 | 92.9 | 29.6 | 85 | 68.7 | 96.3 |
| *X. campestris* pv. azadirachtae LMG 543 | 89.1 | 98.8 | 49.4 | 92.9 | 29.4 | 84.9 | 67.8 | 96.3 |
| *X. axonopodis* pv. bauhinae LMG 548 | 89.1 | 98.8 | 49.5 | 92.9 | 29.6 | 85.0 | 68 | 96.2 |
| *X. campestris* pv. leana LMG 9048 | 88.7 | 98.7 | 49.5 | 92.9 | 29.4 | 84.9 | 68 | 96.3 |
| *X. campestris* pv. thespesiae LMG 9057 | 88.7 | 98.8 | 49.5 | 92.8 | 29.4 | 85 | 67.9 | 96.3 |
| *X. campestris* pv. vitiscarnosae LMG 939 | 88.7 | 98.7 | 49.5 | 92.9 | 29.4 | 84.7 | 68.2 | 96.3 |
| *X. campestris* pv. vitistrifoliae LMG 940 | 88.6 | 98.6 | 49.4 | 92.9 | 29.4 | 84.9 | 67.7 | 96.3 |
| *X. axonopodis* pv. martyniicola LMG 9049 | 88.6 | 98.8 | 49.5 | 93 | 29.4 | 84.8 | 68.2 | 96.3 |
| *X. campestris* pv. vitiswoodrowii LMG 954 | 88.2 | 98.7 | 49.3 | 92.9 | 29.4 | 85 | 68.1 | 96.2 |
| *Pseudomonas cissicola* LMG 21719 | 86.9 | 98.4 | 49.3 | 92.8 | 29.6 | 85.1 | 67.6 | 96.1 |
| *X. axonopodis* pv. khayae LMG 753 | 86.3 | 98.5 | 49.6 | 92.9 | 29.5 | 84.9 | 68.8 | 96.4 |
| *X. campestris* pv. *trichodesmae* NCPPB 585 | 68.3 | 96.1 | 49.7 | 93 | 29.5 | 84.9 | 89.7 | 98.8 |
| *X. campestris* pv. *merremiae* NCPPB 3114 | 67.2 | 96.2 | 49.6 | 92.9 | 29.4 | 85.0 | 88.5 | 98.7 |
| *X. campestris* pv. *thirumalacharii* NCPPB 1452 | 68.3 | 96.2 | 49.7 | 93 | 29.6 | 84.9 | 88 | 98.7 |
| *X. axonopodis* pv. *sesbaniae* NCPPB 582 | 68.2 | 96.1 | 49.7 | 92.9 | 29.6 | 84.9 | 89.6 | 98.8 |
| *X. axonopodis* DSM 3585 (T) | 50.3 | 93.1 | 100 | 100 | 29.3 | 84.9 | 50.2 | 93.1 |
| *X. campestris* pv. *campestris* ATCC 33913 (T) | 30 | 85.1 | 29.3 | 84.9 | 100 | 100 | 29.9 | 85 |
| *X. citri* pv. *fuscan* LMG 726 (T) | 68.6 | 96.3 | 50.2 | 93.1 | 29.9 | 85 | 100 | 100 |

In order to infer the phylogeny of reference pathovars with *X. citri, X. axonopodis* and *X. campestris* we have performed multi locus sequence type (MLST) and core genome-based phylogeny. For MLST six genes namely *fusA, gltA, gapA, gyrB, lacF, lepA* (http://www.pamdb.org) were used and their multiple sequence alignment was generated using clustalw (Thompson, Higgins et al. 1994). Mega7 was used to generate he phylogenetic based tree using neighbor joining (NJ) method with 500 bootstrap (figure 1 A). Further, core genome based phylogenetic tree was generated using roary v3.11.2 (Page, Cummins et al. 2015) with an identity cutoff of 95 (figure 1 B). MLST and whole genome-based phylogeny clearly depicted that the reference pathovars in the present study are related to *X. citri*.

**Figure 1:**
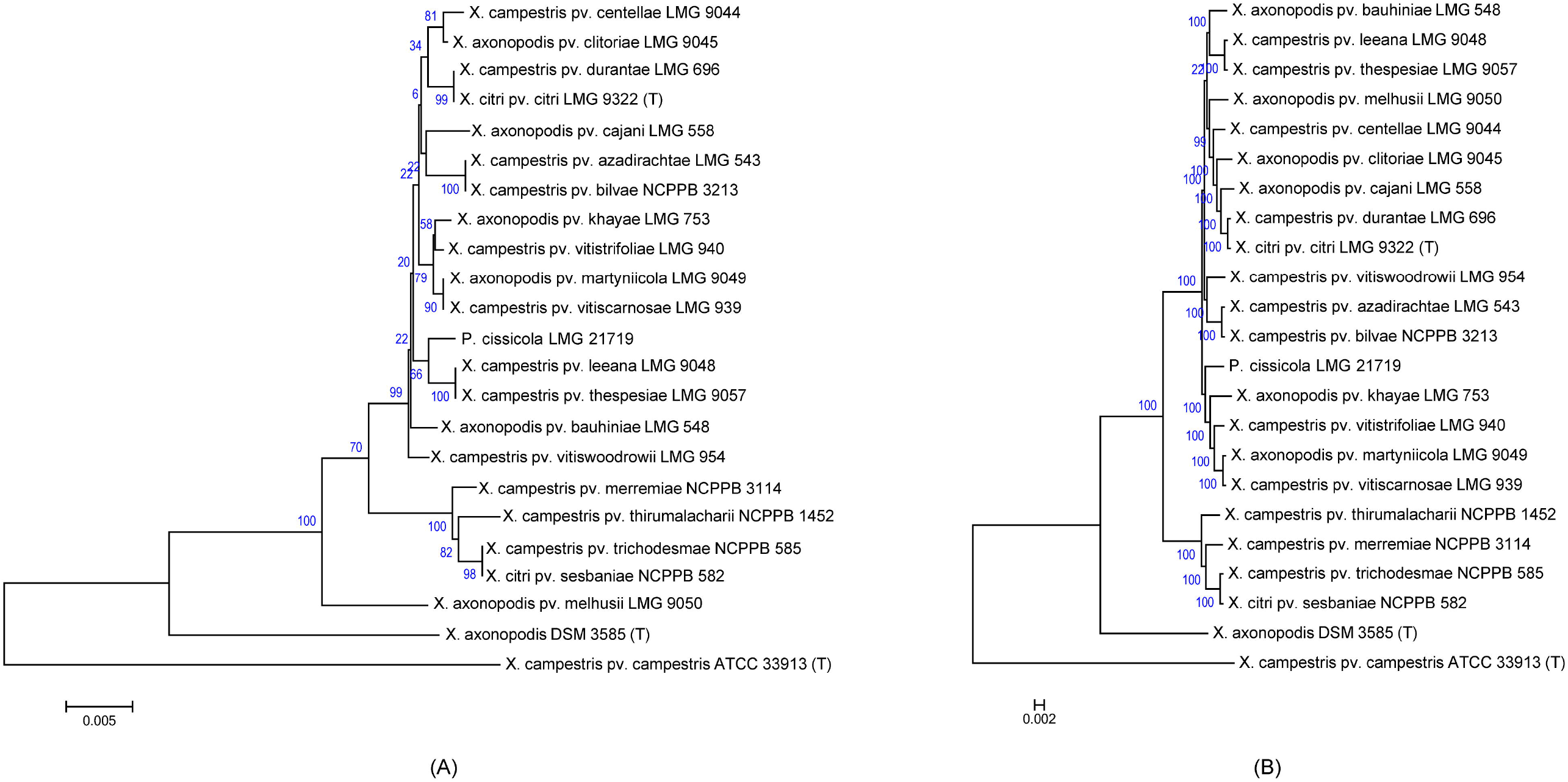
Phylogenetic tree. A). Phylogenetic tree obtained using concatenated MLST gene (*fusA, gltA, gapA, gyrB, lacF, lepA*). B). Phylogenetic tree obtained using the core gene obtained using roary.

Aim of the present study is to emend the taxonomic status of these pathovars and to formally transfer them to the *X. citri*. Owing to the quarantine status of the *X. citri* pv. citri along with other economically important pathovars in the XCPs, their correct taxonomy will aid in their correct identification and understanding the evolution of these closely related pathovars and also the species *X. citri*.

## Emended description of *X. citri* (Hasse 1916) comb. nov

The characteristics of the genus and species of *Xanthomonas citri* are as previously described (Vauterin, Hoste et al. 1995, Constantin, Cleenwerck et al. 2016). In addition to the pathovars previously described (Constantin, Cleenwerck et al. 2016) here we are providing MLSA, phylogenomic and taxonogenomic evidences for other pathovars to constitute *X. citri*.

## Emended description of *X. citri* pv. vitiswoodrowii (Patel and Kulkarni 1951a) comb. nov

= *X. campestris* pv. vitiswoodrowii (Patel and Kulkarni 1951a)

Description as provided in Vauterin *et al*. (1995) and extended genome-based evidences in the present study.

Pathotype strain: LMG 954; ATCC 11636; Dye MW1; ICMP 3965; ICPB PV103; NCPPB 1014; Patel 19; PDDCC 3965; VdM 217

## Emended description of *X. citri* pv. bauhiniae (Padhya, Patel et al. 1965a) comb. nov

**=** *X. axonopodis* pv. bauhiniae (Padhya, Patel et al. 1965a)

Description as provided in Vauterin *et al*. (1995) and extended genome-based evidences in the present study.

Pathotype strain: LMG 548; ICMP 5720; NCPPB 1335; PDDCC 5720; VdM 21 t1,t2

## Emended description of *X. citri* pv. martyniicola (Moniz and Patel 1958) comb. nov

**=** *X. axonopodis* pv. martyniicola (Moniz and Patel 1958)

Description as provided in Vauterin *et al*. (1995) and extended genome-based evidences in the present study.

Pathotype strain: LMG 9049; Dye XG1; ICMP 82; NCPPB 1148

## Emended description of *X. citri* pv. vitiscarnosae (Moniz and Patel 1958) comb. nov

**=** *X. campestris* pv. vitiscarnosae (Moniz and Patel 1958)

Description as provided in Vauterin *et al*. (1995) and extended genome-based evidences in the present study.

Pathotype strain: LMG 939; Dye XH1; ICMP 90; NCPPB 1149; PDDCC 90; VdM 140

## Emended description of *X. citri* pv. vitistrifoliae (Padhya, Patel et al. 1965b) comb. nov

**=** *X. campestris* pv. vitistrifoliae (Padhya, Patel et al. 1965b)

Description as provided in Vauterin *et al*. (1995) and extended genome-based evidences in the present study.

Pathotype strain: LMG 940; ICMP 5761; NCPPB 1451; PDDCC 5761; VdM 141

## Emended description of *X. citri* pv. khayae (Sabet 1959) comb. nov

**=** *X. axonopodis* pv. khayae (Sabet 1959)

Description as provided in Vauterin *et al*. (1995) and extended genome-based evidences in the present study.

Pathotype strain: LMG 753; Dye XK1; ICMP 671; NCPPB 536; PDDCC 671; Sabet Kh1; VdM 81

## Emended description of *X. citri* pv. cissicola (Takimoto 1939) comb. nov

**=** *Pseudomonas cissicola* (Takimoto 1939)

Description as provided (Hu, Young et al. 1997) and extended genome-based evidences in the present study.

Pathotype strain: LMG 21719; ATCC 33616; CCM 2888; CCUG 18839; CFBP 2432; CIP 106723; Goto PC1; ICMP 4289; ICMP 8561; JCM 13362; LMG 2167; NCPPB 2982; PDDCC 4289

## Emended description of *X. citri* pv. melhusii (Patel, Kulkarni et al. 1952b) comb. nov

**=** *X. axonopodis* pv. melhusii (Patel, Kulkarni et al. 1952b)

Description as provided in Vauterin *et al*. (1995) and extended genome-based evidences in the present study.

Pathotype strain: LMG 9050; ATCC 11644; Dye XM1; ICMP 619; ICPB XM107; NCPPB 994; Patel 14

## Emended description of *X. citri* pv. bilvae (Chakravarti, Sarma et al. 1984) comb. nov

**=** *X. campestris* pv. bilvae (Chakravarti, Sarma et al. 1984)

Description as provided in Vauterin *et al*. (1995) and extended genome-based evidences in the present study.

Pathotype strain: NCPPB 3213; ICMP 8918.

## Emended description of *X. citri* pv. azadirachtae (Desai, Gandhi et al. 1966) comb. nov

**=** *X. campestris* pv. azadirachtae (Desai, Gandhi et al. 1966)

Description as provided in Vauterin *et al*. (1995) and extended genome-based evidences in the present study.

Pathotype strain: LMG 543; Dye NR1; ICMP 3102; NCPPB 2388; Patel PA; PDDCC 3102; VdM 18

## Emended description of *X. citri* pv. durantae (Srinivasan and Patel 1957) comb. nov

**=** *X. campestris* pv. durantae (Srinivasan and Patel 1957)

Description as provided in Vauterin *et al*. (1995) and extended genome-based evidences in the present study.

Pathotype strain: LMG 696; ICMP 5728; NCPPB 1456; PDDCC 5728; VdM 50

## Emended description of *X. citri* pv. cajani (Kulkarni 1950) comb. nov

**=** *X. axonopodis* pv. cajani (Kulkarni 1950)

Description as provided in Vauterin *et al*. (1995) and extended genome-based evidences in the present study.

Pathotype strain: LMG 558; ATCC 11639; Dye YR1; ICMP 444; ICPB XC111; NCPPB 573; Patel 5; PDDCC 444; VdM 27

## Emended description of *X. citri* pv. clitoriae (Pandit and Kulkarni 1979) comb. nov

**=** *X. axonopodis* pv. clitoriae (Pandit and Kulkarni 1979)

Description as provided in Vauterin *et al*. (1995) and extended genome-based evidences in the present study.

Pathotype strain: LMG 9045; Bradbury B7090; ICMP 6574; IMI B7090; ITCC 2237; NCPPB 3092

## Emended description of *X. citri* pv. centellae (Basnyat and Kulkarni 1979) comb. nov

**=** *X. campestris* pv. centellae (Basnyat and Kulkarni 1979)

Description as provided in Vauterin *et al*. (1995) and extended genome-based evidences in the present study.

Pathotype strain: LMG 9044; ICMP 6746; ITCC P32; NCPPB 3245

## Emended description of *X. citri* pv. thespesiae (Patil and Kulkarni 1981) comb. nov

**=** *X. campestris* pv. thespesiae (Patil and Kulkarni 1981)

Description as provided in Vauterin *et al*. (1995) and extended genome-based evidences in the present study.

Pathotype strain: LMG 9057; ICMP 7466; ITCC P33

## Emended description of *X. citri* pv. leeana (Patel and Kotasthane 1969a) comb. nov

**=** *X. campestris* pv. leeana (Patel and Kotasthane 1969a)

Description as provided in Vauterin *et al*. (1995) and extended genome-based evidences in the present study.

Pathotype strain: LMG 9048; ICMP 5738; ICPB XL107; NCPPB 2229

## Emended description of *X. citri* pv. sesbaniae (Patel, Kulkarni et al. 1952) comb. nov

**=** *X. axonopodis* pv. sesbaniae (Patel, Kulkarni et al. 1952)

Description as provided in Vauterin *et al*. (1995), Constantin *et al*. (2016) and extended genome-based evidences in the present study.

Pathotype strain: NCPPB 582; ATCC 11675; ICMP 367; LMG 867

## Emended description of *X. citri* pv. merremiae (Pant and Kulkarni 1978) comb. nov

**=** *X. campestris* pv. merremiae (Pant and Kulkarni 1978)

Description as provided in (Pant and Kulkarni 1978) and extended genome-based evidences in the present study.

Pathotype strain: NCPPB 3114; ICMP 6747; LMG 9051

## Emended description of *X. citri* pv. thirumalacharii (Padhya and Patel 1964) comb. nov

**=** *X. campestris* pv. thirumalacharii (Padhya and Patel 1964)

Description as provided in Vauterin *et al*. (1995), Constantin *et al*. (2016) and extended genome-based evidences in the present study.

Pathotype strain: NCPPB 1452; ATCC 11675; ICMP 367; LMG 867

## Emended description of *X. citri* pv. trichodesmae (Patel, Kulkarni et al. 1952b) comb. nov

**=** *X. campestris* pv. trichodesmae (Patel, Kulkarni et al. 1952b)

Description as provided in Vauterin *et al*. (1995) and extended genome-based evidences in the present study.

Pathotype strain: NCPPB 585; ATCC 11678; ICMP 5754; LMG 874

## References

Ah-You, N., et al. (2009). “Polyphasic characterization of xanthomonads pathogenic to members of the Anacardiaceae and their relatedness to species of Xanthomonas.” International Journal of Systematic Evolutionary Microbiology 59(2): 306–318.

Bansal, K., et al. (2019). “Xanthomonas sontii sp. nov., a non-pathogenic bacterium isolated from healthy basmati rice (Oryza sativa) seeds from India.” bioRxiv: 738047.

Bansal, K., et al. (2020). “Deep phylo-taxono-genomics (DEEPT genomics) reveals misclassification of Xanthomonas species complexes into Xylella, Stenotrophomonas and Pseudoxanthomonas.” bioRxiv.

Bansal, K., et al. (2020). “Phylogenomic Insights into Diversity and Evolution of Nonpathogenic Xanthomonas Strains Associated with Citrus.” Msphere 5(2): e00087–00020.

Bansal, K., et al. (2019). “Ecological and evolutionary insights into pathogenic and non-pathogenic rice associated Xanthomonas.” bioRxiv: 453373.

Bansal, K., et al. (2017). “Ecological and evolutionary insights into Xanthomonas citri pathovar diversity.” Applied environmental microbiology 83(9): e02993–02916.

Basnyat, S. R. and Y. S. Kulkarni (1979). “New bacterial leafspot of Centella asiatica L. Urban.” Biovigyanam 5: 179–180.

Brunings, A. M. and D. W. Gabriel (2003). “Xanthomonas citri: breaking the surface.” Molecular plant pathology 4(3): 141–157.

Chakravarti, B. P., et al. (1984). “A bacterial leaf spot of bael (Aegle marmelos Correa) in Rajasthan and a revived name of the bacterium.” Current Science 53(09): 488.

Constantin, E., et al. (2016). “Genetic characterization of strains named as Xanthomonas axonopodis pv. dieffenbachiae leads to a taxonomic revision of the X. axonopodis species complex.” Plant Pathology 65(5): 792–806.

Desai, S. G., et al. (1966). “A new bacterial leaf-spot and blight of Azadirachtaindica.” Indian Phytopathology 19: 322–323.

Gabriel, D., et al. (1989). “Reinstatement of Xanthomonas citri (ex Hasse) and X. phaseoli (ex Smith) to species and reclassification of all X. campestris pv. citri strains.” International Journal of Systematic Evolutionary Microbiology 39(1): 14–22.

Gottwald, T. R., et al. (2002). “Citrus canker: the pathogen and its impact.” Plant Health Progress 3(1): 15.

Gottwald, T. R. and M. Irey (2007). “Post-hurricane analysis of citrus canker II: predictive model estimation of disease spread and area potentially impacted by various eradication protocols following catastrophic weather events.” Plant Health Progress 8(1): 22.

Graham, J. H., et al. (2004). “Xanthomonas axonopodis pv. citri: factors affecting successful eradication of citrus canker.” Molecular plant pathology 5(1): 1–15.

Hasse, C. H. (1916). Pseudomonas Citri, the Cause of Citrus Canker, JSTOR.

Hauben, L., et al. (1997). “Comparison of 16S ribosomal DNA sequences of all Xanthomonas species.” International Journal of Systematic Evolutionary Microbiology 47(2): 328–335.

Hayward, A. (1993). The hosts of Xanthomonas. Xanthomonas, Springer: 1–119.

Hu, F.-P., et al. (1997). “Transfer of Pseudomonas cissicola (Takimoto 1939) Burkholder 1948 to the genus Xanthomonas.” International Journal of Systematic Evolutionary Microbiology 47(1): 228–230.

Kulkarni, Y. S. P. M.K.; Abhyankar, S.G. (1950). “A new bacterial leaf-spot and stem canker of pigeon pea.” Current Science 19(12): 384.

Kumar, S., et al. (2019). “Phylogenomics insights into order and families of Lysobacterales.” Access microbiology 1(2).

Moniz, L. and M. K. Patel (1958). “Three new bacterial diseases of plants from Bombay State.” Current Science 27(12): 494–495.

Moore, E. R., et al. (1997). “16S rRNA gene sequence analyses and inter-and intrageneric relationships of Xanthomonas species and Stenotrophomonas maltophilia.” FEMS microbiology letters 151(2): 145–153.

Naushad, S., et al. (2015). “A phylogenomic and molecular marker based taxonomic framework for the order Xanthomonadales: proposal to transfer the families Algiphilaceae and Solimonadaceae to the order Nevskiales ord. nov. and to create a new family within the order Xanthomonadales, the family Rhodanobacteraceae fam. nov., containing the genus Rhodanobacter and its closest relatives.” Antonie van Leeuwenhoek 107(2): 467–485.

Padhya, A. and M. Patel (1964). “BACTERIAL LEAF-SPOT ON TRIUMFETTA PILOSA ROTH.” Current Science 33(11): 342–342.

Padhya, A. C., et al. (1965a). “A new bacterial leaf-spot disease of Bauhinia racemosa Lamk.” Current Science 34(7): 224–225.

Padhya, A. C., et al. (1965b). “A new bacterial leaf-spot on Vitistrifolia.” Current Science 34(15): 462–463.

Page, A. J., et al. (2015). “Roary: rapid large-scale prokaryote pan genome analysis.” Bioinformatics 31(22): 3691–3693.

Pandit, V. M. and Y. S. Kulkarni (1979). “Bacterial leaf-spot of ClitoriabifloraDalz.” Biovigyanam 5: 9–20.

Pant, N. and Y. Kulkarni (1978). “Bacterial leaf spot of Merremia gangetica (L.) Cufod.” Biovigyanam.

Parkinson, N., et al. (2007). “Phylogenetic analysis of Xanthomonas species by comparison of partial gyrase B gene sequences.” International Journal of Systematic Evolutionary Microbiology 57(12): 2881–2887.

Parkinson, N., et al. (2009). “Phylogenetic structure of Xanthomonas determined by comparison of gyrB sequences.” International Journal of Systematic Evolutionary Microbiology 59(2): 264–274.

Patel, A. M. and W. V. Kotasthane (1969a). “Bacterial blight of Leeaedgeworthii incited by Xanthomonasleeanum, nov. sp.” Current Science 38(21): 519–520.

Patel, M., et al. (1952). “Two new bacterial diseases of plants.” Current Science 21(3): 74–75.

Patel, M. K. and Y. S. Kulkarni (1951a). “A New Bacterial Leaf Spot on Vitis Woodrowil Stapf.” Current Science 20(5): 132.

Patel, M. K., et al. (1952b). “Some new bacterial diseases of plants.” Current Science 21(12): 345–346.

Patil, A. S. and Y. S. Kulkarni (1981). “A new bacterial leaf-spot disease of Thespesiapopulnea Sol. ex Corr.” Current Science 50(23): 1040–1041.

Pruvost, O. (1993). “Xanthomonas campestris pv. mangiferaeindicae: cause of bacterial black spot of mangoes.”

Pruvost, O., et al. (1998). “Phenotypic diversity of Xanthomonas sp. mangiferaeindicae.”

Raychaudhuri, S., et al. (1972). “The history of plant pathology in India.” Annual Review of Phytopathology 10(1): 21–36.

Sabet, K. A. (1959). “Studies in the bacterial diseases of Sudan crops IV. Bacterial leaf-spot and canker disease of mahogany (Khayasenegalensis (Desr.) A. Juss. and K. grandifoliola C. DC).” Annals of Applied Biology 47: 658–665.

Schaad, N. W., et al. (2006). “Emended classification of xanthomonad pathogens on citrus.” Papers in Plant Pathology: 96.

Srinivasan, M. C. and M. K. Patel (1957). “Two new phytopathogenic bacteria on verbenaceous hosts.” Current Science 26(03): 90–91.

Takimoto, S. (1939). “Bacterial leaf spot of Cissus japonica Willd.” Annals of the Phytopathological Society 9: 41–43.

Thompson, J. D., et al. (1994). “CLUSTAL W: improving the sensitivity of progressive multiple sequence alignment through sequence weighting, position-specific gap penalties and weight matrix choice.” Nucleic acids research 22(22): 4673–4680.

Vauterin, L., et al. (1995). “Reclassification of xanthomonas.” International Journal of Systematic Evolutionary Microbiology 45(3): 472–489.

Vauterin, L., et al. (2000). “Synopsis on the taxonomy of the genus Xanthomonas.” Phytopathology 90(7): 677–682.

Vauterin, L., et al. (1996). “Identification of non-pathogenic Xanthomonas strains associated with plants.” Systematic applied microbiology 19(1): 96–105.

Yoon, S.-H., et al. (2017). “A large-scale evaluation of algorithms to calculate average nucleotide identity.” Antonie van Leeuwenhoek 110(10): 1281–1286.

